# SCHEEPDOG: programming electric cues to dynamically herd large-scale cell migration

**DOI:** 10.1101/2019.12.20.884510

**Authors:** Tom J. Zajdel, Gawoon Shim, Linus Wang, Alejandro Rossello-Martinez, Daniel J. Cohen

## Abstract

Directed cell migration is critical across biological processes spanning healing to cancer invasion, yet no existing tools allow real-time interactive guidance over such migration. We present a new bioreactor that harnesses electrotaxis—directed cell migration along electric field gradients—by integrating four independent electrodes under computer control to dynamically program electric field patterns, and hence steer cell migration. Using this platform, we programmed and characterized multiple precise, two-dimensional collective migration maneuvers in renal epithelia and primary skin keratinocyte ensembles. First, we demonstrated on-demand, 90-degree collective turning. Next, we developed a universal electrical stimulation scheme capable of programming arbitrary 2D migration maneuvers such as precise angular turns and migration in a complete circle. Our stimulation scheme proves that cells effectively time-average electric field cues, helping to elucidate the transduction time scales in electrotaxis. Together, this work represents an enabling platform for controlling cell migration with broad utility across many cell types.

## Introduction

As directed and large-scale collective cell migration underlie key multicellular processes spanning morphogenesis, healing, and cancer progression, a tool to shepherd such migration would enable new possibilities across cell biology and biomedical engineering (Friedl and Gilmour, 2009). Such a tool must be: (1) broadly applicable across multiple cell and tissue types, and (2) programmable to allow spatiotemporal control. Chemotaxis, an obvious candidate, lacks broad applicability as it requires ligand-receptor interactions. Furthermore, diffusion causes difficulties with interactive, spatiotemporal chemical dosing (Kress *et al*., 2009; Li and Lin, 2011; Berthier and Beebe, 2014). Alternately, micropatterned proteins or topographies can direct motion across cell types, but cannot be dynamically changed (Xiong *et al*., 2019). Finally, optogenetics requires genetic modification and must be targeted to individual cells at a subcellular level to guide migration (Weitzman and Hahn, 2014).

Mounting evidence suggests electrical cues as the ideal basis for general cellular herding. Since du Bois-Reymond’s discovery 175 years ago that skin wounds possess an endogenous electric field(Du Bois-Reymond, 1849), it is now well-established that: (1) 1-5 V cm^-1^ electric fields are natural responses to ionic imbalances generated in vivo during morphogenesis, regeneration, and pathogenesis (Nuccitelli, 2003; McCaig, Song and Rajnicek, 2009); (2) cells transduce DC electrical cues into navigational cues and migrate along the field gradient in a process called ‘electrotaxis’ or ‘galvanotaxis’; and (3) electrotaxis is remarkably conserved across at least 20 diverse mammalian cell types, cellular slime molds, zebrafish, and frogs (McCaig, Song and Rajnicek, 2009; Cortese *et al*., 2014). Recent work has revealed that endogenous electric fields likely act on membrane-bound receptors via electrophoresis (electrostatic force) or electro-osmotic-flow (shear force) rather than directly affecting intracellular components (Allen, Mogilner and Theriot, 2013). While transduction remains an active research area in electrotaxis and bioelectricity as a whole, it is exciting to note that electrotaxis and chemotaxis share key overlapping signaling via TORC2 and PI3K/PTEN, which modulate front-rear polarity (Zhao *et al*., 2006; Zhao, Devreotes and VanHook, 2015). Hence, electrotaxis is a powerful phenomenon that should allow us to electrically program key migratory pathways, giving it unique potential as a tool for controlling cell migration.

As electric fields can be harnessed to direct cellular migration, modern electronic tools and approaches should enable unprecedented control over collective cell dynamics. However, contemporary electrotaxis work emphasizes electrotaxis as a phenomenon rather than a tool, and engineering and technology approaches have received less attention. Nearly all electrotactic devices use static DC fields with single anode/cathode pairs (Tai *et al*., 2009; Tandon *et al*., 2009; Daniel J. Cohen, Nelson and Maharbiz, 2014; Sroka *et al*., 2018; Gokoffski *et al*., 2019), restricting migration control to fixed, 1D trajectories. The potential of 2D stimulation is not realized by the handful of existing dual-axis studies due to lack of programmable control of field direction (Gokoffski *et al*., 2019), and production of cytotoxic byproducts (Riedel-Kruse *et al*., 2011). Further, the vast majority of electrotaxis studies focus on single cells due to experimental complexities, while the smaller body of collective-level studies (Cooper and Keller, 1984; Nishimura, Isseroff and Nuccitelli, 1996; Zhao *et al*., 1996; Cao, Pu and Zhao, 2011; Li *et al*., 2012; Daniel J. Cohen, Nelson and Maharbiz, 2014; Lalli and Asthagiri, 2015; Bashirzadeh *et al*., 2018; Cho *et al*., 2018; Gokoffski *et al*., 2019) focused on specific biological or biophysical questions rather than pushing the envelope of electrotaxis to explore both its limits and capabilities as a tool. A next-gen electro-bioreactor is key to pushing electrotaxis research forward by enabling large-scale, precise, and repeatable perturbations for cell motility research. It would also offer new ways to control tissues and morphogenesis for tissue engineering and regenerative medicine, where there is already considerable interest in bioelectricity to manipulate developmental processes and expedite wound healing; both processes that use endogenously generated electric fields as part of their signaling pathways (Levin, Pezzulo and Finkelstein, 2017; Long *et al*., 2018; Mathews and Levin, 2018; Kriegman *et al*., 2020).

To illustrate this potential we present the only instance of freely programmable, 2D herding of collective cell migration, enabled by our next-gen electro-bioreactor called SCHEEPDOG: Spatiotemporal Cellular HErding with Electrochemical Potentials to Dynamically Orient Galvanotaxis. Here, we describe the design of the SCHEEPDOG system, and then validate it by programming millimeter-scale, on-demand collective maneuvers in renal epithelia and primary skin cell ensembles numbering over 10,000 individuals. Finally, we demonstrate a universal migratory control framework using 4 interactive electrodes to take advantage of, and shed light on, the key timescales involved in electrotaxis. These findings highlight the profound plasticity of collective migration and emphasize that electrotaxis may well be the most broadly applicable, programmable migratory cue for herding cell migration in real-time.

## Results

### Design of next-gen electro-bioreactor for multi-axis migration control

Developing electrotaxis into a viable tool to programmatically herd cells requires a fundamentally new approach to bioelectric stimulation: interactive, spatiotemporal control of the electric field, and thereby of migration. SCHEEPDOG (Fig. 1) accomplishes this with three key design modules: (1) bioreactor architecture; (2) life support; and (3) dynamically programmable electric field generation via 4 independent electrodes.

**Figure 1.**
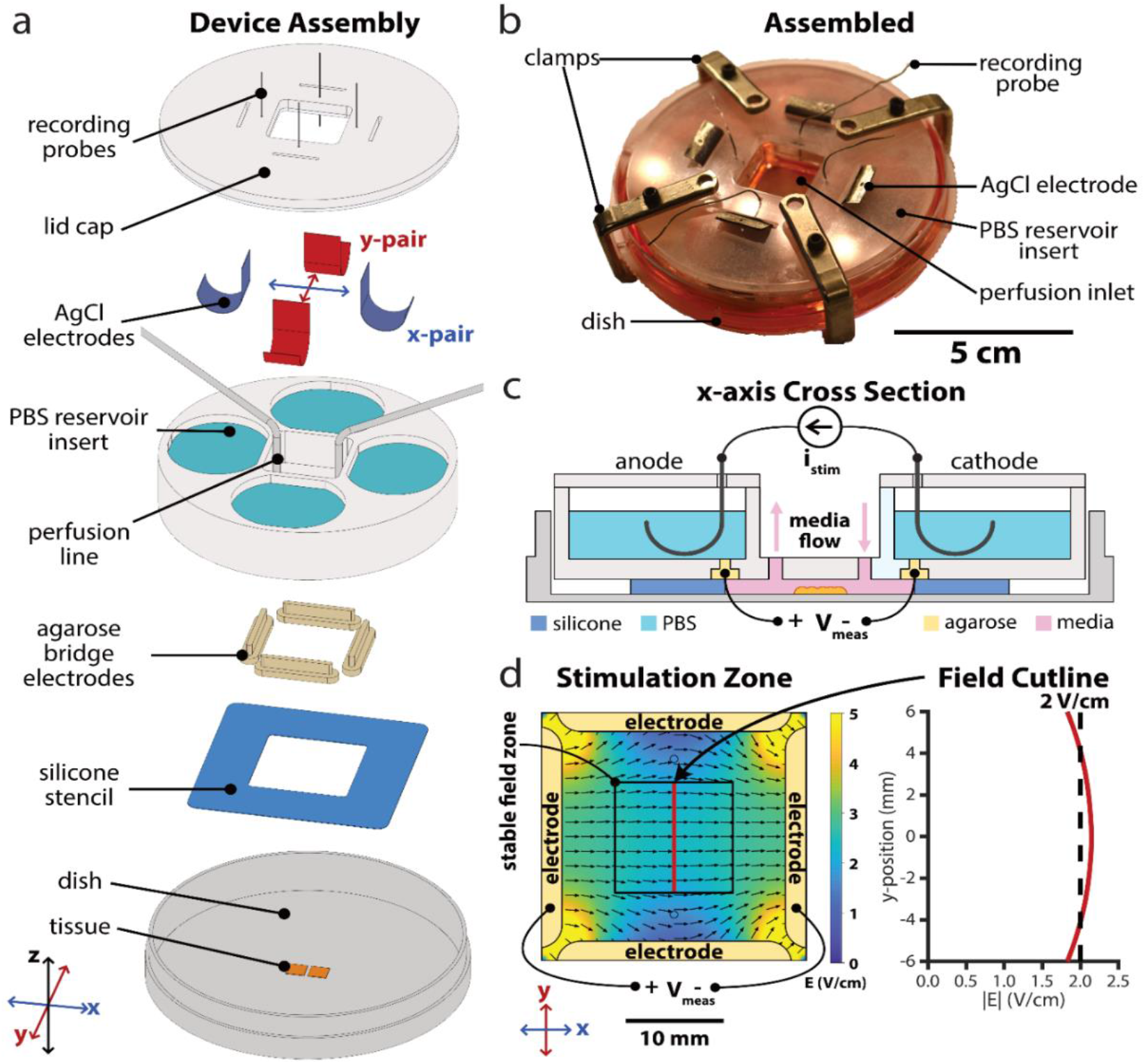
Design of next-gen electro-bioreactor for multi-axis migration control; SCHEEPDOG: Spatiotemporal Cellular HErding with Electrochemical Potentials to Dynamically Orient Galvanotaxis, a bioreactor for two-dimensional, bioelectric control over collective migration. (a) Assembly of the SCHEEPDOG bioreactor within a Petri dish. A silicone stencil defines a stimulation zone in the center of the dish, and tissues are patterned within this region. Four agarose bridges held inside a laser-cut PBS reservoir insert are then pressed against the stencil to fully enclose the stimulation zone. Ag/AgCl electrodes and perfusion lines are added, then a lid is clamped to the assembly and recording probes are inserted into each agarose bridge electrode. (b) Photograph of a fully assembled device. (c) Sectional schematic of one axis of SCHEEPDOG. Each axis features one anode/cathode pair connected to a computer-controlled current supply which provides istim. A digital oscilloscope records channel voltage Vmeas to enable closed-loop control. (d) Numeric simulation of the field generated by the device when the horizontal electrode pair is stimulated. The simulated field value is indicated with a red line showing a stable field zone in the center of the device. See also Supplementary Figs. 1,2.

The bioreactor housing uses rapid, in situ microfluidic assembly around pre-patterned cells or tissues, and integrates best-practices from our prior work and that of others (Daniel J Cohen, Nelson and Maharbiz, 2014; Sun, 2017; Cho *et al*., 2018) (Fig. 1, Supplementary Fig. 1). It also ensures stability and consistency by reducing variable conditions in previous assembly procedures of microfluidic devices, many of which rely on vacuum grease and glass coverslip-ceilings (Cao, Pu and Zhao, 2011; Gokoffski *et al*., 2019). Briefly, cells or tissues are patterned on a substrate using a silicone seeding stencil to define the tissue shape, and then a separate silicone foundation layer is sandwiched between a laser-cut insert and the cell culture substrate to define a stimulation zone (Fig. 1d). Confining cells to a narrow stimulation zone concentrates DC electric fields and extends electrode lifespan (Daniel J Cohen, Nelson and Maharbiz, 2014). Further, such in situ assembly affords complete control of cell and tissue geometry and composition, as well as compatibility with arbitrary culture substrates such as large glass coverslips.

To accommodate metabolic needs and prevent electrochemical cytotoxicity, we integrated several life-support and safeguard systems into SCHEEPDOG. Virtually all DC electrotaxis systems inject Faradaic current across the cells through an anode and a cathode, typically comprised of silver chloride or noble metals immersed in saline reservoirs (Song *et al*., 2007). SCHEEPDOG used two pairs of Ag/AgCl electrode foils in saline reservoirs organized along two orthogonal axes, each independently connected to a dedicated computer-controlled current source. This compensates for typical electrochemical variations during stimulation such as potential drift of Ag/AgCl electrodes as they are consumed during stimulation. The silver foil electrodes were chloridized electrochemically (Methods), (Vulto *et al*., 2009; Huang *et al*., 2013) to improve electrode longevity over common, bleach-based methods. To prevent cytotoxic electrochemical byproducts produced at each electrode-electrolyte interface from reaching cells, most electrotaxis systems integrate a salt-bridge diffusion barrier (Schopf, Boehler and Asplund, 2016) while others use continuous media perfusion across the cells (Cole and Gagnon, 2019). SCHEEPDOG integrated both structured agarose bridges (Huang *et al*., 2013; Song *et al*., 2013; Hou *et al*., 2014; Wu *et al*., 2015) and continuous media perfusion to keep tissues oxygenated and free of electrochemical byproducts (Methods). These measures contribute to SCHEEPDOG’s exceptionally long, stable run-times (8-12 h at least) without any detectable cytotoxicity despite the high driving current (typically 8-10 mA).

The heart of SCHEEPDOG is interactive, spatiotemporal control of the electric field via 4 stimulation electrodes grouped into orthogonal pairs (e.g. X and Y axes). Because incorporating two anode-cathode pairs required a large stimulation region, special care was taken to ensure field uniformity to improve throughput by designing the integrated agarose bridges to occupy nearly the entire perimeter of the stimulation chamber (Huang *et al*., 2001; Daniel J Cohen, Nelson and Maharbiz, 2014; Tsai *et al*., 2016). Current leak through the unstimulated pair was also minimized by using narrow agarose bridges rather than a plus-shaped chamber, thus improving field uniformity relative to prior efforts (Supplementary Fig. 2). To ensure a temporally stable field throughout extended experiments, SCHEEPDOG used closed-loop feedback and monitored the channel voltage with a pair of probing electrodes (Fig. 1c). To ensure uniform stimulation, tissues were kept within a 12×12 mm square region in the center of the stimulation chamber where field uniformity varied by only ±7% across the culture zone according to the numerical simulation of the field in COMSOL (Fig. 1d). Such precision control allowed us to freely program complex stimulation patterns while maintaining constant stimulation strength and field uniformity (Methods).

### Validating 2-axis control of collective cell migration

We selected a 90° turn as an archetypal complex maneuver to validate bi-axial, programmable control over collective cell migration. To capture a range of phenotypes, we tested with both the MDCK kidney epithelial cell line and primary, neonatal mouse skin keratinocytes. While both cell lines undergo electrotaxis in 1D (Nishimura, Isseroff and Nuccitelli, 1996; Li *et al*., 2012; Daniel J Cohen, Nelson and Maharbiz, 2014; Guo *et al*., 2015; Bashirzadeh *et al*., 2018), primary keratinocytes cultured in basal media have weak cell-cell adhesions leading to more individualistic behavior in counterpoint to the strong cadherin-mediated cell-cell adhesions in an MDCK epithelium.(Li *et al*., 2012) We patterned large, 5×5 mm monolayers of either cell type into SCHEEPDOG, then used automated phase-contrast microscopy to capture 1 h of control data (field OFF) before stimulating at 2 V cm^-1^ for 2 h along the X-axis (‘right’) followed by 2 h along the Y-axis (‘up’). Initially, we tracked ~2000 cells from a 2×2 mm central zone of each monolayer to avoid edge effects and overlaid these trajectories to reflect the ensemble migration responses (Figs. 2a,d; Supplementary Videos 1,2). Analysis shows that each population clearly underwent a 90° turn as seen in Figs. 2a,d where the mean trajectories are indicated by solid white (‘right’) and black (‘up’) lines.

To better quantify these maneuvers, we analyzed migration in the bulk using tools from swarm dynamics and collective cell migration (Poujade *et al*., 2007; Daniel J Cohen, Nelson and Maharbiz, 2014). Using particle image velocimetry (PIV), we generated migration vector flow fields across the tissues at each time point (Methods), and used these vector fields to calculate the directionality order parameter, *ϕ*, which reflects how well cells obeyed the ‘right’ and ‘up’ commands. *ϕ* is defined as the average of the cosine or sine of every PIV vector with respect to the direction of the electrical command, as shown in Equations 1 and 2:

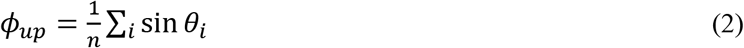

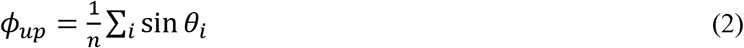

*ϕ* can vary between −1 (anti-parallel) to 1 (perfect alignment), and Figs. 2b,d depict *ϕ_right_* and *ϕ_up_* throughout programmed 90° turns. Both cell types exhibited rapid response to the initial ‘rightward’ command, reaching peak *ϕ_right_* > 0.9, and similar performance for *ϕ_up_*. However, when the command switched from ‘right’ to ‘up’, *ϕ_right_* was quite slow to decrease back to baseline, indicating persistent rightward migration after the ‘up’ command had been given, with *ϕ_right_* for MDCKs taking ~90 min to return to baseline, while keratinocytes took closer to ~60 min.

**Figure 2.**
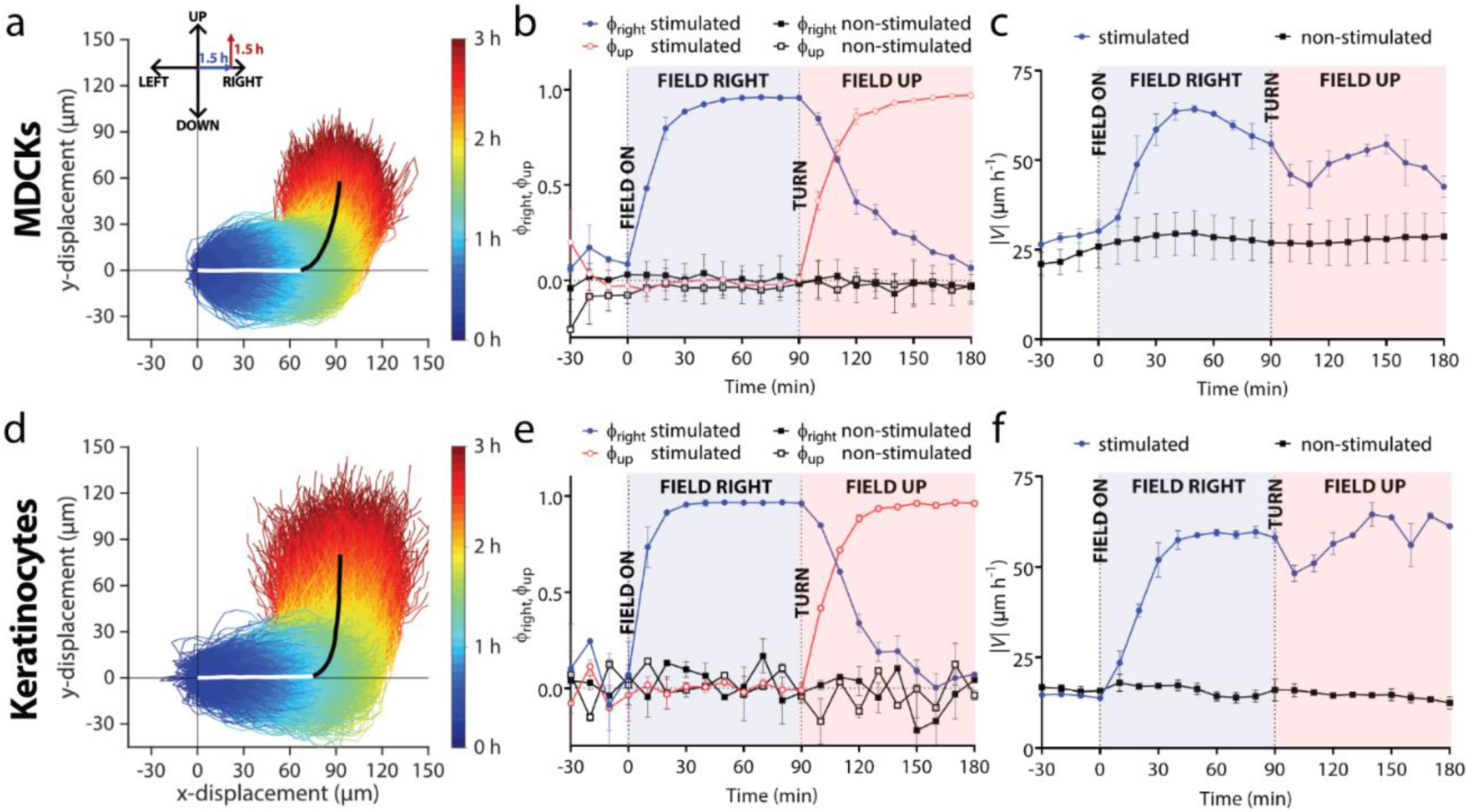
Validating 2-axis control of collective cell migration with a demonstration of collective 90° turns in monolayers of either MDCK cells or keratinocytes (trajectories measured from n = 2 monolayers of each cell type). (a,d) Temporal color-coded trajectories of individually tracked MDCKs (a, n = 4,086 trajectories) or keratinocytes (d, n = 2,642 trajectories) stimulated for 1.5 h ‘right’, and then 1.5 h ‘up’. For visual clarity, trajectories within one standard deviation of the mean trajectory (bold white and black overlaid lines indicating the two phases of the turn) are selected and plotted.. Individual trajectories were taken from 2×2 mm central segments of 5×5 mm tissues during induced migration. (b,e) Mean directional order parameter both in the horizontal (blue) and vertical (red) axis during a turn for MDCKs (b) or keratinocytes (e). Data are plotted in 10 min intervals and represent the PIV-derived order averaged across tissues (n = 2). Error bars represent standard deviation across tissues. Dashed vertical lines denote when the field was switched on or changed direction. (c,f) Mean velocity in the analysis region for MDCKs (c) or keratinocytes (f). Error bars represent the standard deviation across tissues.

To a large ensemble collectively moving rightwards, a sudden command to make a 90° turn is a strong perturbation that should affect migration speed. We quantified this using the PIV data (Figs. 2c,f) and observed first a characteristic rise in migratory speed upon initiating electrotaxis that largely persisted for the duration of the experiment. However, we also noticed distinct, transient dips in speed immediately following the command switching from ‘right’ to ‘up’ for each cell type. While the trajectories appear quite smooth (Figs. 2a,d), there was clearly an overall slow-down during the turning maneuver as cells reorient and realign with their neighbors, likely reflecting shifting traction forces and general neighbor-neighbor collisions (Daniel J Cohen, Nelson and Maharbiz, 2014; Cho *et al*., 2018). Once the turn was complete, speed began to rise again for both cell types as cells entrained to the new direction.

While MDCKs exhibited a much smoother response with a tighter trajectory spread, keratinocytes tracked the signal more accurately at an ensemble level (Figs. 2a,b). MDCK monolayers exhibited considerable mechanical memory manifesting as persistent rightward motion even as the tissues turned upwards. By contrast, keratinocyte ensembles turned more sharply with less apparent memory of the previous migration direction. When we quantified this by characterizing the overshoot and time for a given population to complete a turn (Supplementary Fig. 3), we observed that keratinocytes reoriented nearly 30% more quickly than MDKCs. This prompted us to further explore the ‘controllability’ of tissues using SCHEEPDOG to better understand the limitations of electrotactic maneuvers.

### Programming large-scale tissue translation

An exciting aspect of programming directed cell migration is the capability to physically translate or steer the growth of a tissue, which has important implications spanning collective migration research, tissue engineering, and wound healing. Surprisingly, we observed radically different boundary-level control responses between MDCK and keratinocyte ensembles despite both exhibiting qualitatively similar bulk responses. To explore this, we analyzed how cells in different parts of the population—the trailing and leading edges vs. the central bulk—responded during the first 90 min phase of the turning maneuver. Strikingly, keratinocytes exhibited nearly uniform guided migration across the entire ensemble (50-70 μm translation per cell), while MDCK monolayers experienced almost no directed migration at the leading or trailing edges in contrast to the strong directed response in the bulk (5 μm vs. 50+ μm) as can be seen in Fig. 3a. The difference between the two cell types in their boundary dynamics during electrotaxis is emphasized in Fig. 3b where kymographs show leading edge translation. Such a location-dependent cellular response in MDCK monolayers (edge vs. bulk) suggests a supracellular response, likely due to pronounced cell-cell adhesion (Daniel J Cohen, Nelson and Maharbiz, 2014; Cho *et al*., 2018; Shellard *et al*., 2018; Alert and Trepat, 2020), while keratinocyte ensembles migrate more akin to ‘marching in formation’ as a choreographed unit; exhibiting much tighter, population-wide responses.

**Figure 3.**
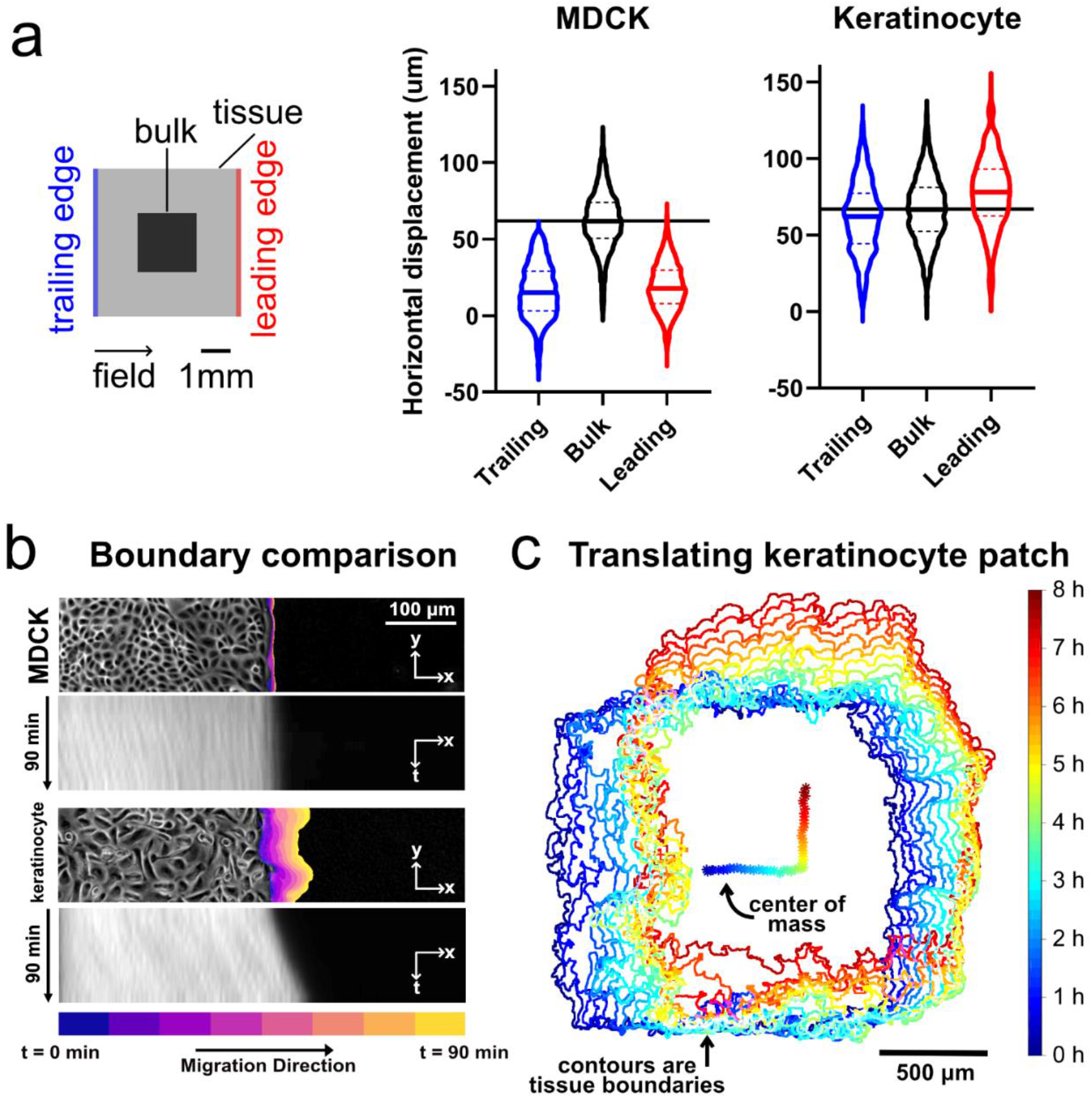
Comparing edge and bulk behavior during electrotaxis. We compared MDCK and keratinocyte monolayer electrotaxis during 90 min of single-axis stimulation. The electric field was directed ‘right’ for 1.5 h, then ‘up’ for 1.5 h. Data were analyzed from the 2×2 mm central segments of 5×5 mm tissues. (a) Visualization of the mean trajectory overshoot from the ideal trajectory (green) of the MDCK (blue) and keratinocyte (red) tissues. Trajectories averaged across n = 2 tissues for each cell type (4,086 MDCK trajectories and 2,642 keratinocyte trajectories). (b) Comparison of the overshoot, response time, relaxation time and turn time. Data plotted are the same as from panel (a), with least-squares fit lines overlaid. (c) Boundary displacement and kymographs for MDCK and keratinocyte tissues during the 90 min stimulation to the right. Boundary displacement plots visually indicate extent of outgrowth while the kymographs depict the average edge outgrowth across the entire leading edge of representative tissues. (d) Boundary displacement and tracking of the ensemble center of mass of a keratinocyte monolayer over an 8 h run, with 4 h stimulation in the ‘right’ and 4 h ‘up’. See also Supplementary Videos 1-3.

Taking advantage of large-scale keratinocyte displacement, we pushed the limits of coordinated migration by programming an 8 h turning maneuver with 4 h of stimulation ‘rightward’ and ‘upward’, respectively into a small keratinocyte ensemble (Supplementary Video 3). Fig. 3c presents both the tissue boundary and the trajectory of the tissue centroid which clearly traces out a sharp 90° turn with monolayer translation of up to 500 μm (1/3 of the group size) in each direction over time. While a small fraction (~10%) of the population lagged behind, suggesting heterogeneity in the electrotactic response, the overall ensemble response was remarkably stable and the initial square formation was largely preserved (Supplementary Video 3). Overall, these data demonstrate that extraordinary displacements can be programmed as long as the native collective dynamics of the target cell-type are taken into account.

### Developing a universal migration control scheme

A truly programmable migration controller ought to support arbitrary maneuvers. To achieve this with our 4-electrode design, we hypothesized that switching between orthogonal X- and Y-field stimulation faster than the electrotactic transduction time would make cells perceive the time-averaged direction and migrate in that direction. Here, any desired 2D maneuver can be represented as a serialized sequence of independent X- or Y-axis migration command pulses, akin to an ‘etch-a-sketch’ drawing. In this control scheme, we fixed the field strength at ~2 V cm^-1^, and then altered only the relative duration of X-axis commands versus Y-axis commands, as shown in Fig. 4a.

**Figure 4.**
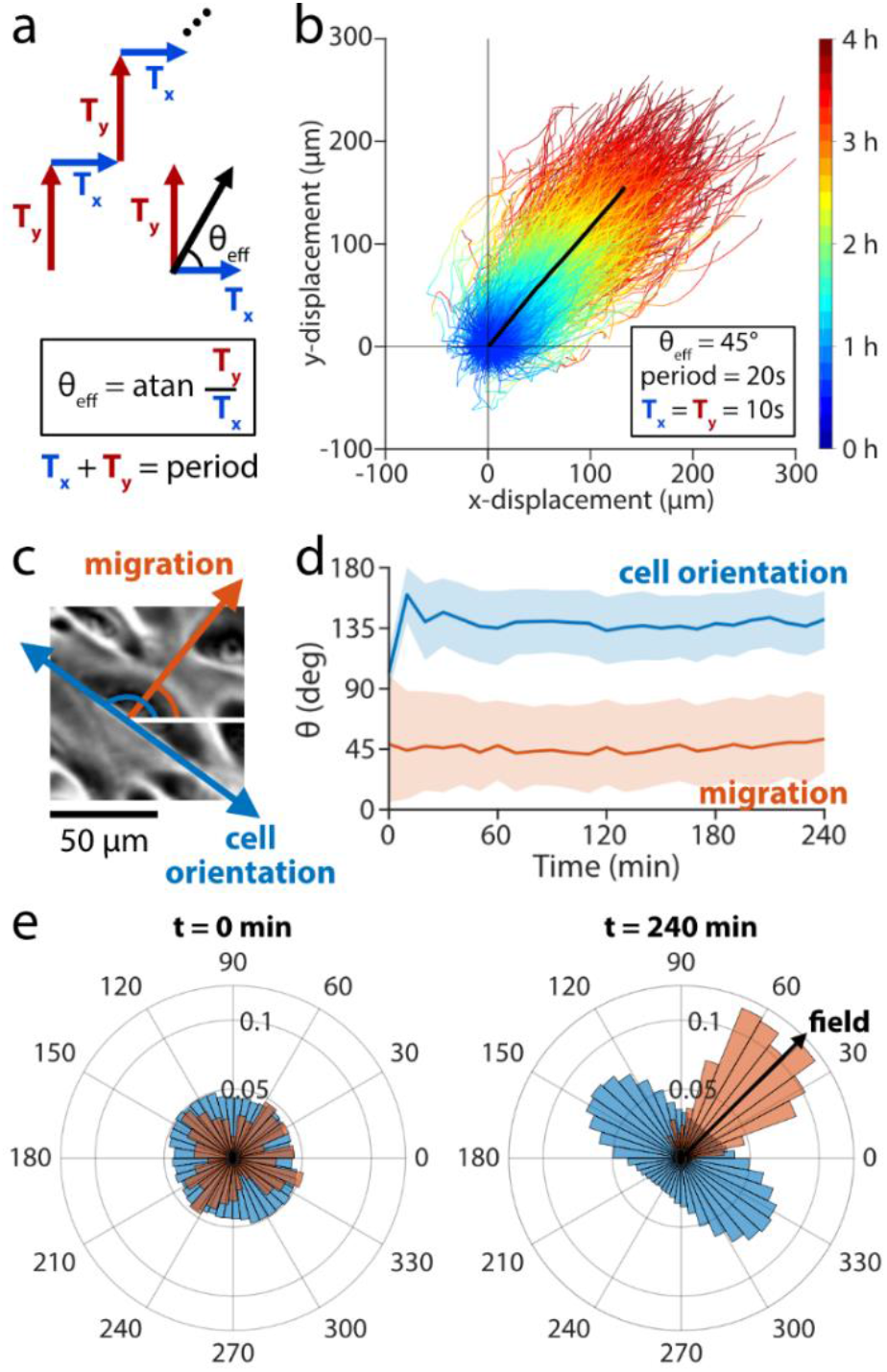
Developing a universal migration control scheme based on rapid switching of electric field directions to enable arbitrary field angles. (a) The stimulation sequence divides a 20 sec stimulation window into two periods: horizontal stimulation (*T_y_*) and vertical stimulation ( *T_y_*). The ratio of *T_y_/T_x_* determines the effective stimulation angle that the cells experience. (b) Trajectories of 2×2 mm central segments from keratinocyte ensembles across 4 h of stimulation with θeff = 45° (n = 2 ensembles). The mean trajectory is plotted as a thick black line and followed a migration angle of 44°. Tracks within two standard deviations of the mean cell trajectory are selected and plotted as time-coded multicolor lines (n = 900 trajectories). (c) Phase contrast image of keratinocyte cell migrating at ~45° (red arrow), showing the cell body oriented along ~135° (blue arrow). (d) Dominant orientation angle as computed by Fourier components analysis (blue) shows a rapid orientation to an average of 135° ± 20° throughout stimulation. Migration angle (red) averages 44° ± 32° throughout stimulation. Shaded bars represent standard deviation in orientation across one representative ensemble. (e) Cell body orientation (blue) and migration direction (red) histograms for a representative monolayer at t = 0 min, just as stimulation began, and at t = 240 min, just as stimulation ended. Cell body orientation ranges from 0° to 180° and its distribution is duplicated across the horizontal for visualization purposes. Cells in this representative monolayer averaged an orientation of 136° ± 20° and migration direction of 50° ± 28° just as stimulation ended. The cells across the ensemble oriented perpendicularly to the migration direction of 45°. See also Supplementary Videos 4,5.

If this framework were valid, a symmetric stimulation sequence (‘right’, ‘up’, repeat) would induce a 45° migration trajectory without any ‘wobble’ if we stimulated quickly enough. We tested a *T_y_/T_x_* ratio of 1, where each direction was alternately stimulated for 10 sec (50% duty cycle) to produce a 45° effective angle with the horizontal (Fig. 4a,b; Supplementary Video 4). Over a total duration of 4 h, the monolayer displaced 147 μm in the horizontal direction and 143 μm in the vertical, tracking a ~45° angle overall (Supplementary Video 5) and suggesting that the method of ‘virtual angle’ control is effective within ~30° for the majority of cells over any 10 min period. Notably, the long axis of individual cell bodies aligned orthogonal to the effective electric field vector, resulting in cell bodies stably orienting at ~135° (Fig. 4c-e), consistent with prior observations in 1D electrotactic systems (Hammerick, Longaker and Prinz, 2010; Cho *et al*., 2018).

That cells migrate smoothly along a 45° trajectory when symmetrically stimulated between ‘right’ and ‘up’ in quick succession proves that cells can time-average electrotactic commands to perceive a virtual command given sufficiently rapid stimulation switching. We also validated the generality of this control scheme by programming a different cell type—MDCKs—to undergo a 60° turn (Supplementary Fig. 4, Supplementary Video 6). Together, these data suggest that the field-sensing mechanism of electrotaxis operates on the timescale of seconds, much faster than the migration response, which occurs on the order of 10 minutes (Sawhney and Howard, 2002; Cheng and Zygourakis, 2007), enabling cells to time-average rapidly varying electrical cues.

### Herding cells through a continuous, arbitrary maneuver

To explore the capabilities and limits of SCHEEPDOG’s control, we programmed a more complex maneuver into a keratinocyte monolayer–a closed circle—that required continuously adjusted stimulation commands. By modulating the ratio *T_x_/T_y_* over time, we discretized the circular path into 80 serial turns of approximately 4.5° equally spaced over an 8 h period (Fig. 5a).

**Figure 5.**
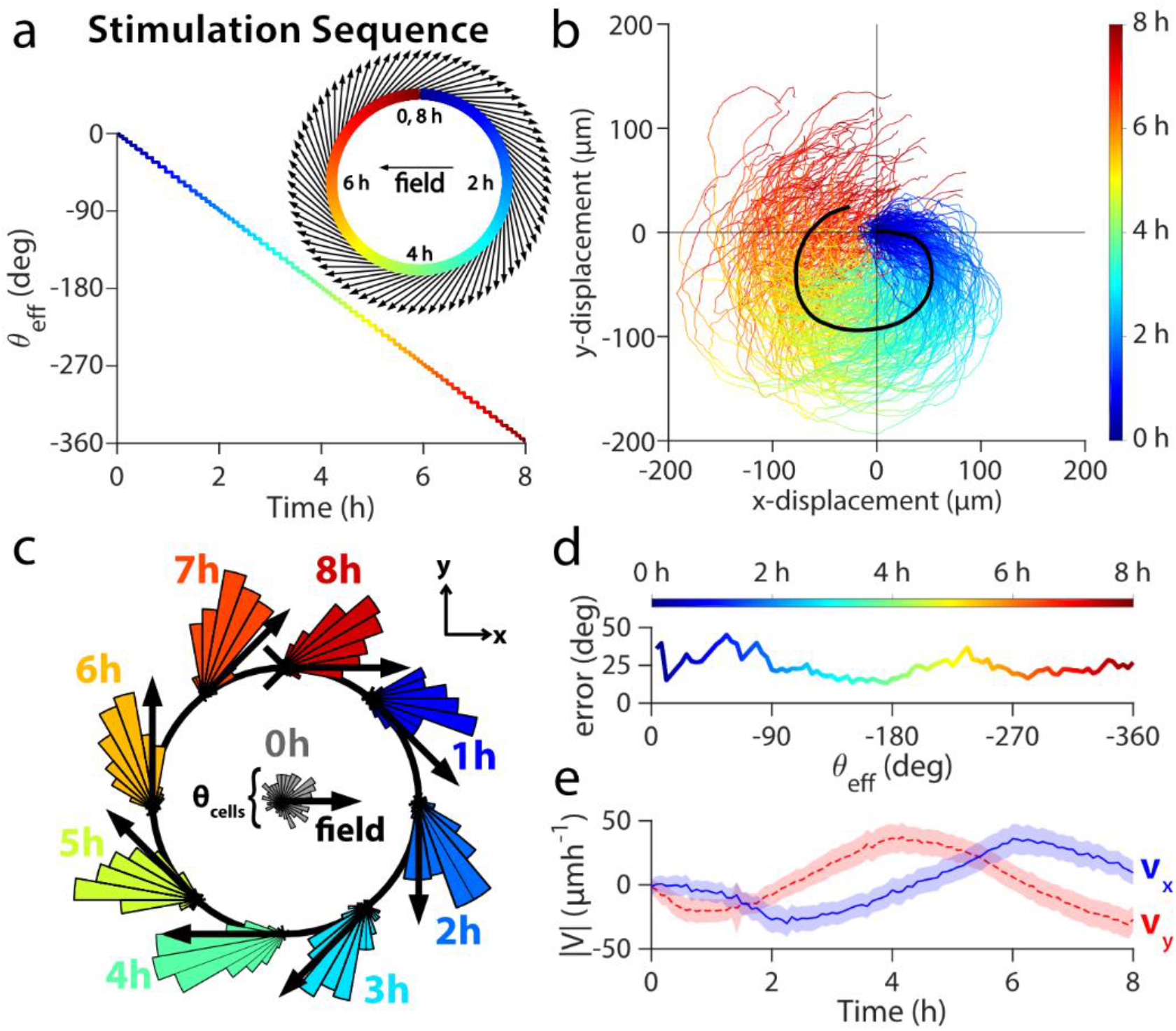
Herding cells in a closed circle to demonstrate general, continuous control. (a) The stimulation sequence used for discretization of the circular maneuver with respect to time. θeff angle across time for the entire circle is plotted as a multicolor line, while the discretization is overlaid as a black line. Insets show a close up of the first hour of θeff over time (upper right) and the effective field vectors of stimulation for this period (lower left). (b) Trajectories of 2×2 mm central segments from two keratinocyte tissues throughout 8 h circular stimulation (n = 224 trajectories). (c) Angular histogram of cell velocities calculated from PIV. Black arrows indicate the direction of stimulation. (d) Phase lag between the mean cell migration direction and the effective stimulation direction θeff. (e) PIV-derived velocity averaged across two tissues in X- (blue) and Y-directions (blue). Shaded region represents standard deviation. See also Supplementary Video 7.

Remarkably, cells successfully tracked this continuously shifting electric field, completing a circular maneuver with an average perimeter of over 300 μm (Fig. 5b; Supplementary Video 7). To quantify the population dynamics during this maneuver, we generated angular histograms of migrating tissues at key time-points throughout the 8 h period (Fig. 5c). These show consistent tracking of the command vector where cells lag behind by an average of 25° over the maneuver (Fig. 5d), suggesting intrinsic temporal limitations of electrotactic maneuvers at the population level. Velocity analysis (Fig. 5e) shows that both vx and vy varied sinusoidally and ~90° out of phase with each other, characteristics expected of circular motion. These data validate that truly arbitrary programming of cellular maneuvers is possible by discretizing complex maneuvers, highlight the remarkable plasticity of ensemble migration patterns, and represent the first example to our knowledge of a tissue obeying a prescribed, continuously varying 2D migrational cue.

## Discussion

SCHEEPDOG’s active, multi-electrode design literally allows large groups of cells to be herded across a wide range of cellular systems and exemplifies the surprising potency and versatility of electrotaxis as both a phenomenon and a tool in its own right. We developed SCHEEPDOG to address the lack of existing tools to program and steer cell migration beyond what is possible with contemporary ‘-taxis’ platforms (Fuller *et al*., 2010), and we ensured that our approach can be manufactured and deployed quickly from biology to engineering labs. The unique integration of 4 addressable electrodes with layer-based microfluidics and bioreactor design elements allows SCHEEPDOG to produce uniform stimulation patterns for extended live-cell experiments. We demonstrated this by programming truly arbitrary migration maneuvers ranging from diagonal lines to full circles using a generalized stimulation scheme that scales to any 2D maneuver by rapidly switching between stimulation electrode pairs.

At a fundamental research level, SCHEEPDOG can be used to elucidate signaling and the biomechanics of electrotaxis. For instance, the data we gathered using SCHEEPDOG’s dynamic stimulation provides insights into the key timescales involved in electrotaxis. Specifically, our results support the hypothesis that there are two key timescales governing electrotaxis: (1) rapid sensing of the electric field (~10 sec), and (2) slower cellular polarization and migration response (5-10 minutes). This separation of scales also occurs in chemotaxis, which has been shown to share certain overlapping signaling and migration infrastructure with electrotaxis (Gao *et al*., 2015). Our data in Fig. 4_that show keratinocytes moving diagonally at 45° in response to rapid and symmetrically alternating X- and Y-direction stimulation (i.e. every 10 sec) suggest that the cells responded to a time-averaged field direction, and the realignment of cell bodies perpendicular to this axis of translation without a discernible ‘wobble’ further supports our hypothesis that cells sensed a composite field from these orthogonal stimuli. The single, steady-state response generated when the frequency of switching was 0.33Hz implies that this signal was time-averaged by a slower process in the electrotaxis signal transduction pathways operating on a longer timescale. This is consistent with the dominant hypothesis in the electrotaxis field that DC stimulation induces the asymmetric distribution of membrane-bound receptors that leads to front-rear polarization. Prior studies on model receptors indicate that full receptor aggregation occurs on the order of 5 minutes during stimulation at physiological field strengths (typically 1-2 V cm^-1^, though as high as 5 V cm^-1^ in some systems) (Allen, Mogilner and Theriot, 2013; Sarkar *et al*., 2019). As we cycle the stimulation direction between ‘right’ and ‘up’ for 10 seconds in each direction to create a 45° diagonal trajectory, it is unlikely that affected receptors would fully polarize within any individual stimulation period. The time-averaging behavior, therefore, likely stems from gradually increasing polarization of affected receptors throughout the ‘right’/’up’ cycling; the receptors gradually aggregating along the time-averaged axis, to establish front-rear polarity. Such malleable receptor aggregation would be compatible with accounts implicating PI3K, PTEN, and PIP2/3 in the electrotaxis transduction process, much like in chemotaxis (Zhao *et al*., 2006; Meng *et al*., 2011). Future research on the mechanisms of electrotaxis could employ SCHEEPDOG’s dynamic switching capabilities to investigate not only the localization time of common polarity factors, but also the spatiotemporal dynamics of induced cytoskeletal reorganization and genetic sources of heterogeneity in the response to electric cues across individual cells within a monolayer to further our understanding of how the electrical signal is transduced to establish cell polarity and directed motion.

SCHEEPDOG offers new possibilities not only for electrotaxis research in its own right, but also for the broader space of cell migration, tissue engineering, and bioelectricity research. As studies have shown that the vast majority of tested mammalian cell types undergo electrotaxis, SCHEEPDOG will be valuable in studying collective migration for an exceptionally broad range of cell types and applications (Figs. 2,3) (McCaig, Song and Rajnicek, 2009; Cortese *et al*., 2014). The ability to precisely modify geometry and time scales of cell migration in real time can provide unique perturbations tailored for specific assays such as cytoskeletal rearrangement, signaling, transcription, and mechanical processes during directed migration. Further, we foresee this form of interactive cell migration control underlying the development of a variety of ‘bioelectric band-aids’ to guide and accelerate wound healing, angiogenesis, and other morphogenic processes that rely on endogenous DC electric fields (Borgens, Vanable and Jaffe, 1977; Barker, Jaffe and Vanable Jr, 1982; Zhao *et al*., 2004; McLaughlin and Levin, 2018). In addition to these endogenous systems, synthetic biological approaches have generated new electrically-driven transcription circuits that suggest exciting applications for programmable electrical cues (Weber *et al*., 2009). Electrotaxis has also been shown to differentially affect tumor cells, so interactive electrical control might offer approaches to inhibit tumor outgrowth (McCaig, Song and Rajnicek, 2009; Hou *et al*., 2014). Synthetic biological approaches have generated new electrically-driven transcription pathways that could be interactively controlled by SCHEEPDOG. Ultimately, we hope that SCHEEPDOG will help to standardize and refine both design practices and applications of electrotaxis across research fields, promote accessibility and reproducibility, inspire new directions in electrotaxis and biological research, and better allow us to engineer and harness electrotaxis as a powerful and unique sheepdog.

## Methods

### Cell culture

Wild-type MDCK-II cells (courtesy of the Nelson Laboratory, Stanford University) were cultured in Dulbecco’s Modified Eagle’s Medium with 1 g L^-1^ glucose, 1 g L^-1^ sodium bicarbonate, 10% fetal bovine serum, and 1% penicillin–streptomycin. Primary keratinocytes were harvested from mice (courtesy of the Devenport Laboratory, Princeton University) and cultured in E-medium supplemented with 15% serum and 50 μM calcium. All cells were maintained at 37 °C under 5% CO2 and 95% relative humidity. Cells were split before reaching 70% confluence and passage number was kept below 20 for MDCKs and 30 for keratinocytes for all experiments.

### Device fabrication and assembly

SCHEEPDOG was designed for simultaneous fluid perfusion and electrical stimulation. The chamber geometry was designed in CAD software (Autodesk Fusion) and then simulated in finite element software (COMSOL) to fine-tune the resultant electric field pattern (Fig. 1d) and fluid perfusion dynamics. The chamber pattern was cut from a 250 μm thick sheet of silicone rubber (Bisco HT-6240, Stockwell Elastomers) by a computer-controlled cutter (Cameo, Silhouette). This silicone stencil was applied to the center of a 10 cm diameter plastic tissue culture dish (Falcon). This stencil formed the watertight outline of the electro-stimulation zone against the plastic substrate. In the case of keratinocytes, fibronectin was adsorbed to the dish’s surface to provide a matrix for cellular adhesion (protein dissolved to 50 μg mL^-1^ in DI water, applied to the dish for 30 min at 37 °C, then washed with DI water). To seed tissues in the stimulation zone and control the shape and size of monolayers, a second silicone stencil containing square microwells was cut and applied to the center of the culture substrate. With the stencil in place, a seeding solution of cells was prepared at a density of 2.2 × 10^6^ cells for MDCKs and 1.7 × 10^6^ cells for keratinocytes. 10 mL of the cell solution was pipetted into the tissue stencils at a ratio of 2 μL of solution per 5 mm^2^ of stencil area. Extra humidification media was added to the periphery of the tissue culture dish and MDCK cells were allowed to settle for 1 h, whereupon sufficient media was added to fill the dish and it was left to incubate for 16 h. For keratinocytes, the cells were allowed to settle for 6 h and incubated for 14 h to account for substrate adhesion differences.

The complete device assembly consisted of four Ag/AgCl stimulation foil electrodes, four Ti wire recording electrodes, an acrylic salt reservoir insert, and a lid cap. The three acrylic pieces comprising the reusable salt water reservoir insert were all cut from a 5.2 mm thick acrylic sheet using a computer controlled laser cutter (VLS 3.5, Universal Laser Systems). These individual layers of acrylic were stacked, clamped, then solvent welded together with acrylic cement (SCIGRIP 4, SCIGRIP Assembly Adhesives) and left to set for 72 h. The lid cap for the assembly was cut from a 3 mm thick acrylic sheet and a 1 mm-thick silicone sheet was attached to its bottom side to provide a better seal against the reservoir insert. All components were sterilized by exposure to 5 min UV radiation in a cell culture hood before assembly.

After the incubation period, the cell stencil was removed and the device was assembled within 2 h after the stencil removal. To produce the integrated salt bridges, agarose was melted into phosphate buffered saline at pH 7.4 (PBS) at 4% w/v on a hot plate. Once fully melted, the agarose was cast into the four slots in the main insert to serve as bridges. Once the agarose bridges had solidified, 3 mL of PBS was added to each of the salt reservoirs. Then, the filled and cast insert was placed over the tissue culture dish and aligned against the silicone stencil and pressed into place, taking care to avoid trapping air bubbles between the stimulation zone and the insert. Next, the four Ag/AgCl stimulation electrodes were inserted into the lid cap and the lid cap was pressed against the reservoir insert. The four recording electrodes were then pushed through the lid cap, each one making contact with one of the agarose bridges. Once everything was in place, four modified C-clamps (Humboldt Manufacturing Co.) were used to hold the assembly together. Once assembled, the device was immediately brought to the microscope for imaging.

### Electrochemical system

#### Electrode preparation

To produce the stimulation electrodes, 125 μm silver foil (Alfa Aesar) was cut into 1.25 cm by 5 cm strips and chloridized electrochemically; the silver working electrodes were immersed in 0.25 M KCl and subsequently connected to the positive terminal of a programmable DC power supply (Rigol DP832), and a 25 cm coiled Ti wire (0.5 mm, Alfa Aesar) was used as the counter electrode to drive 100 μA cm^-2^ plating current for 8 h. Once plated, the electrodes were washed in DI and slightly curled in preparation for assembly. Four Ti wires (0.5 mm diameter, Alfa Aesar) were cut to 5 cm length to serve as voltage recording electrodes for the closed-loop voltage controller.

#### Instrumentation

Two Keithley source meters (Keithley 2400/2450, Tektronix) provided current to the stimulation electrode pairs, with each meter providing one axis of stimulation. A USB oscilloscope (Analog Discovery 2, Digilent Inc.) measured the voltage across each pair of recording electrodes. A custom MATLAB script was used to command the instruments to drive a set current and measure the resultant voltage across the stimulation chamber, using proportional feedback control to adjust the output current required to maintain the target field strength and direction. Only one axis was activated at a time to maximize field uniformity within the stimulation zone. In this work, the field strength was set at ~2 V cm^-1^. To produce intermediate stimulation directions between the horizontal and vertical axes when required, each 20 sec stimulation cycle was divided between a horizontal and vertical stimulation period (Tx and Ty, respectively) in order. The effective stimulation angle approximated by this stimulation scheme can be calculated by Equation 3:

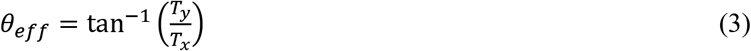

For example, if a 30-degree angle from the horizontal were desired, (*T_x_*, *T_y_*) would be set to (13 sec, 7 sec), so θeff would be 28°. *T_x_* and *T_y_* were programmed to 1 sec resolution, meaning that 20 unique angles were possible over 90 degrees, resulting in an average field direction resolution of 4.5°. Arbitrary stimulation functions, like the 8-hour circle in Fig. 5, were piecewise parameterized to produce a sequence of angles, each lasting 20 sec, which best approximated the function. In practice, the desired angle changed slowly. In the case of the circle, each angle was stimulated for an average of 3 minutes (9 stimulation cycles) before SCHEEPDOG proceeded to the next angle.

### Microscopy

All images were acquired on an automated Zeiss (Observer Z1) inverted fluorescence microscope equipped with an XY motorized stage and controlled using Slidebook (Intelligent Imaging Innovations, 3i). The microscope was fully incubated at 37 °C, and a peristaltic pump (Instech Laboratories) placed within the chamber was used to continually perfuse fresh media through the electro-bioreactor at a rate of 2 mL h^-1^. Media pH was regulated by continuously bubbling 5% CO2 through the inlet media reservoir during perfusion. All imaging was performed using a 5X/0.16 phasecontrast objective. Nuclear labeling was performed using Hoechst 33342 (Invitrogen), a DAPI filter set (358 nm excitation, 461 nm emission), and 30 msec exposure with a metal halide lamp (xCite 120, EXFO). Images were taken at either 5 min or 10 min intervals as indicated in the text.

### Image processing and analysis

All post-processing of tissue microscopy images was performed using FIJI. Images were collected sequentially, tiled, and template matched to correct for stage drift prior to being analyzed for either PIV or cellular tracking analysis.

### Trajectory generation via nuclear tracking

Nuclear labeling was performed using either a custom-written machine-learning classifier trained to detect individual nuclei from phase contrast data (all MDCK data) or using Hoechst labeling (all keratinocyte data). Resulting nuclei were tracked using FIJI’s TrackMate plugin set to detect spots via Laplacian of the Gaussian filtering and to link using Linear Motion Tracking (Tinevez *et al*., 2017). Mean population trajectories for a given maneuver were calculated by averaging the X- and Y-coordinates of all tracks at each time point.

### Particle image velocimetry

Tissue migratory flow fields were generated using PIVLab, a MATLAB script performing FFT-based PIV(Thielicke and Stamhuis, 2014). PIV analysis was performed over the central zone of the target tissue (~65% of the total area) to avoid edge artifacts as described earlier(Daniel J Cohen, Nelson and Maharbiz, 2014). Iterative window analysis was performed using first 96×96 pixel windows followed by 48×48 pixel windows, both with 50% step overlap. Vector validation excluded vectors beyond five standard deviations and replaced them with interpolated vectors. The velocity vector fields were then imported into MATLAB to calculate the speed and migration orientation. Rosette plots that show the direction of motion of each cell at a given time point were extracted by calculating the mean angle across the vector field.

### Boundary edge displacement kymographs

Boundary images and kymographs were produced using FIJI. For the kymographs, the X-position of the tissue edge was averaged across the entire length of the tissue using a median averaging algorithm, then stacked temporally. Boundary images were produced by Gaussian blurring each frame in FIJI to create a binary binding box for the cells, then detecting the edges of each of these frames to overlay them temporally. Boundary displacement distance was calculated by measuring the distance between the maximum X-position or Y-position of the leading edge boundary both before and after stimulation.

### Tissue center of mass

Cell nuclei images were median filtered to reduce background noise, then binarized using thresholding in FIJI. The binary images were them imported into MATLAB to find the tissue centroid at each frame by calculating the ensemble average of the centroids of all nuclei within the tissue.

### Orientation analysis

Quantification of cell orientation was performed using FIJI’s directionality analysis plugin. Fourier components analysis was used to determine a histogram for the distribution of angles present in each phase contrast image(Liu, 1991). A Gaussian function was fit to this histogram. The center of this Gaussian was reported as the dominant angle. Migration angles were computed from the tracked trajectories and the mean and standard deviation of these angles were calculated by the MATLAB CircStat toolbox (Berens *et al*., 2009).

## AUTHOR CONTRIBUTIONS

T.J.Z., G.S., A.R., and D.J.C. designed the stimulation device and experiments and T.J.Z., G.S., and L.W. performed the experiments. T.J.Z, G.S., and D.J.C. wrote the manuscript. D.J.C designed and supervised the study.

## DATA AVAILABILITY STATEMENT

The data that supports the findings of this study are available from the corresponding author upon reasonable request.

## CODE AVAILABILITY STATEMENT

Custom code and scripts used for this study are available from the corresponding author upon reasonable request.

