## Supplementary Information for "SCHEEPDOG: programming electric cues to dynamically herd large-scale cell migration"

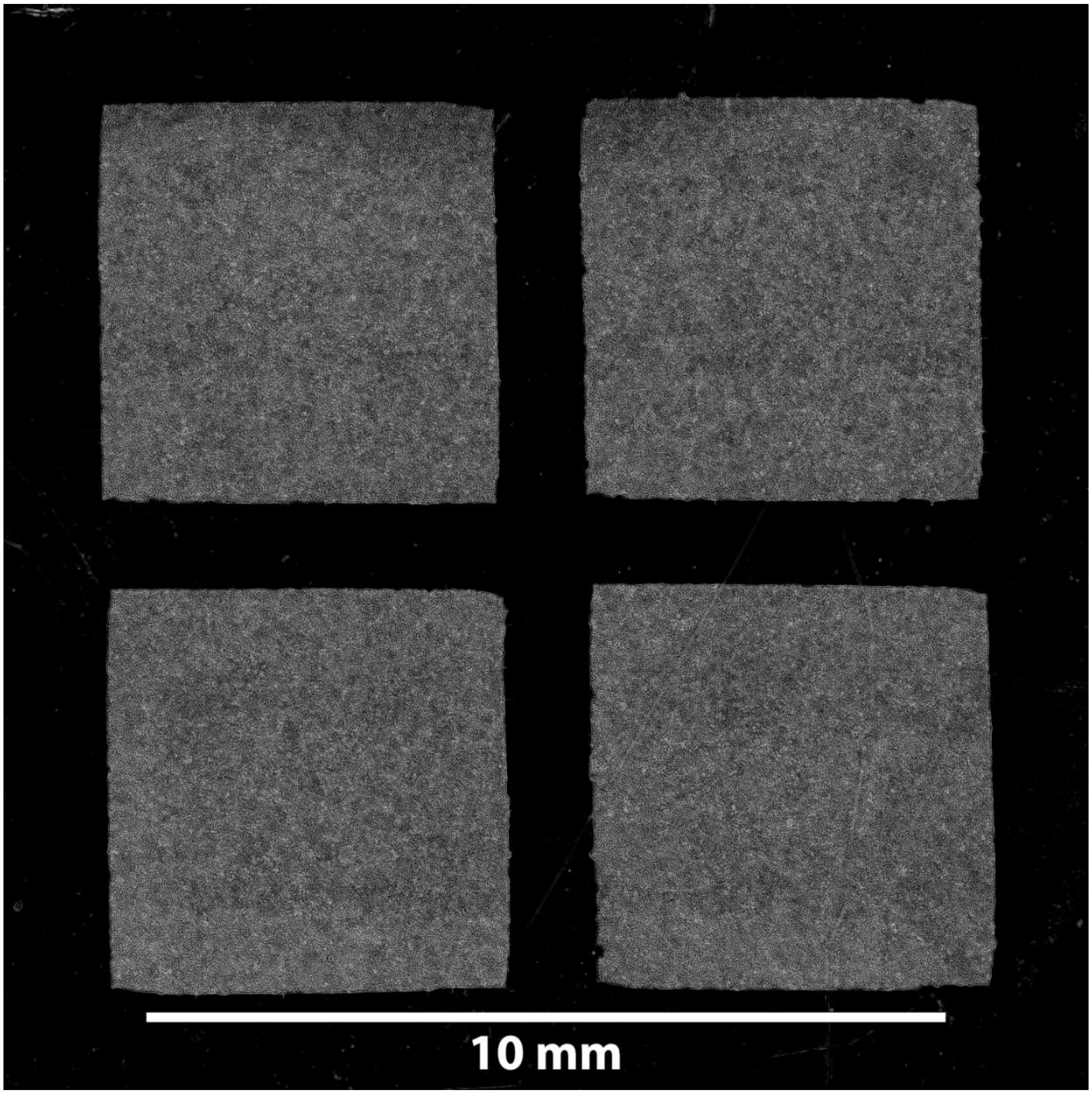


**Supplementary Figure 1:** Phase-contrast microscopy image of four 5×5 mm MDCK monolayers patterned inside the SCHEEPDOG device's stimulation zone. Image was background subtracted and its brightness was increased for visual clarity. Related to Figs. 1,2.

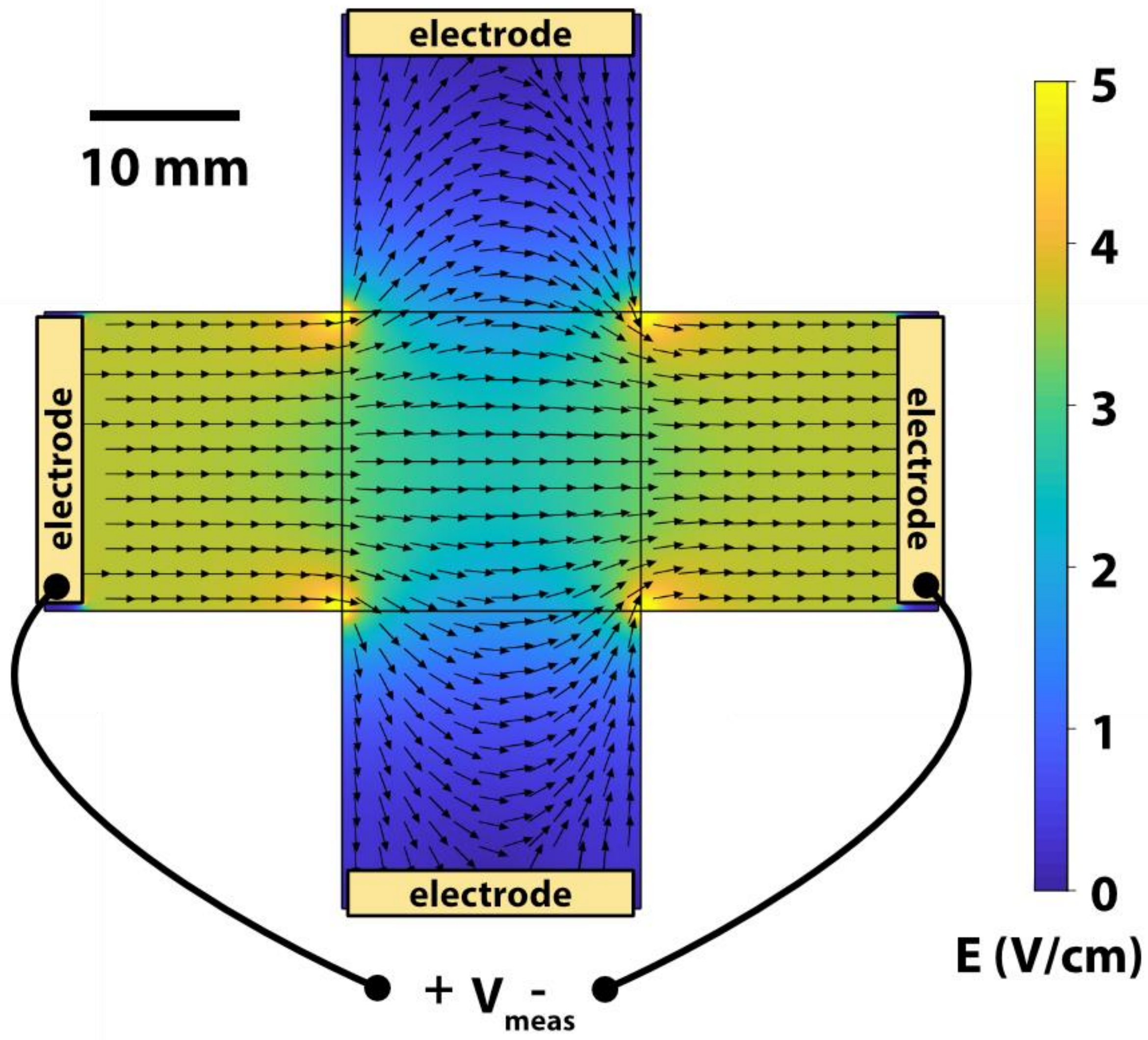


**Supplementary Figure 2:** Simple double-axis chamber design that results in non-uniform field. Care must be taken when adding a second pair of electrodes to an electrochemical stimulation system to avoid excessive current leak through the orthogonal axis. Otherwise, field uniformity in the center is adversely affected. Related to Fig. 1.

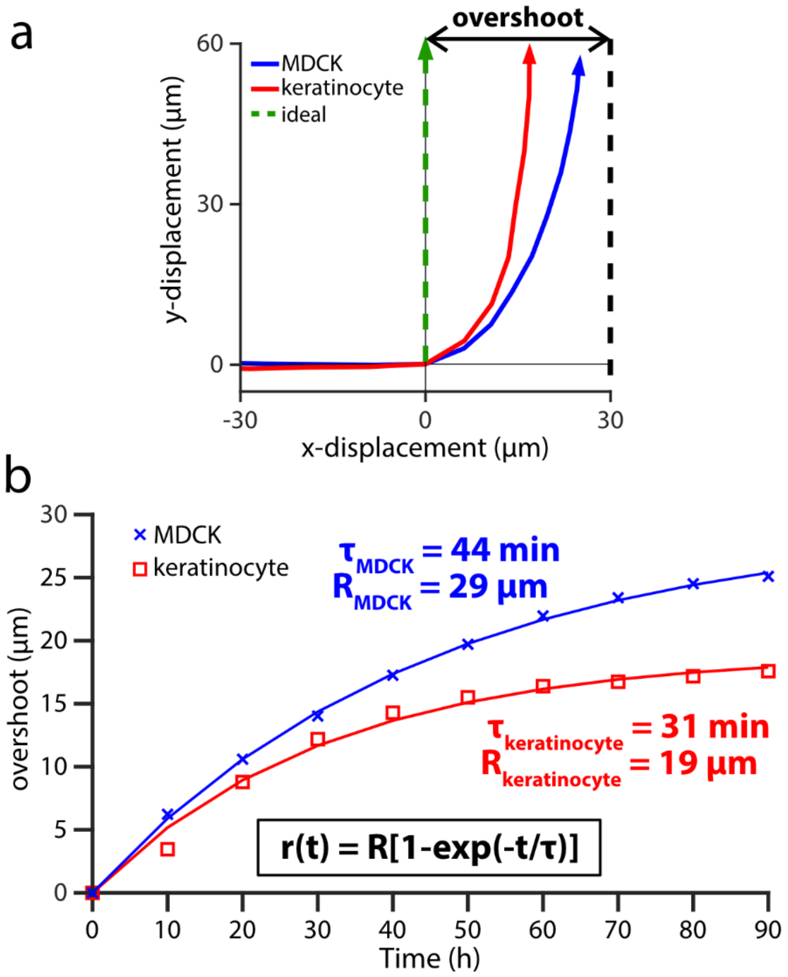


**Supplementary Figure 3:** Defining electrotactic performance metrics. Related to Fig. 3. We compared MDCK and keratinocyte tissue electrotaxis during programmed 90º turns. The electric field was directed 'right' for 1.5 h then 'up' for 1.5 h. Data were analyzed from the 2×2 mm central segments of 5×5 mm tissues. (a) Visualization of the mean trajectory overshoot from the ideal trajectory (green) of the MDCK (blue) and keratinocyte (red) tissues. Trajectories averaged across n = 2 tissues for each cell type (n = 4086 MDCK trajectories and n = 2642 keratinocyte trajectories). (b) Comparison of the overshoot, response time, relaxation time and turn time. Data plotted are the same as from panel (a), with least-squares fit lines overlaid. See also Supplementary Videos 1-3.

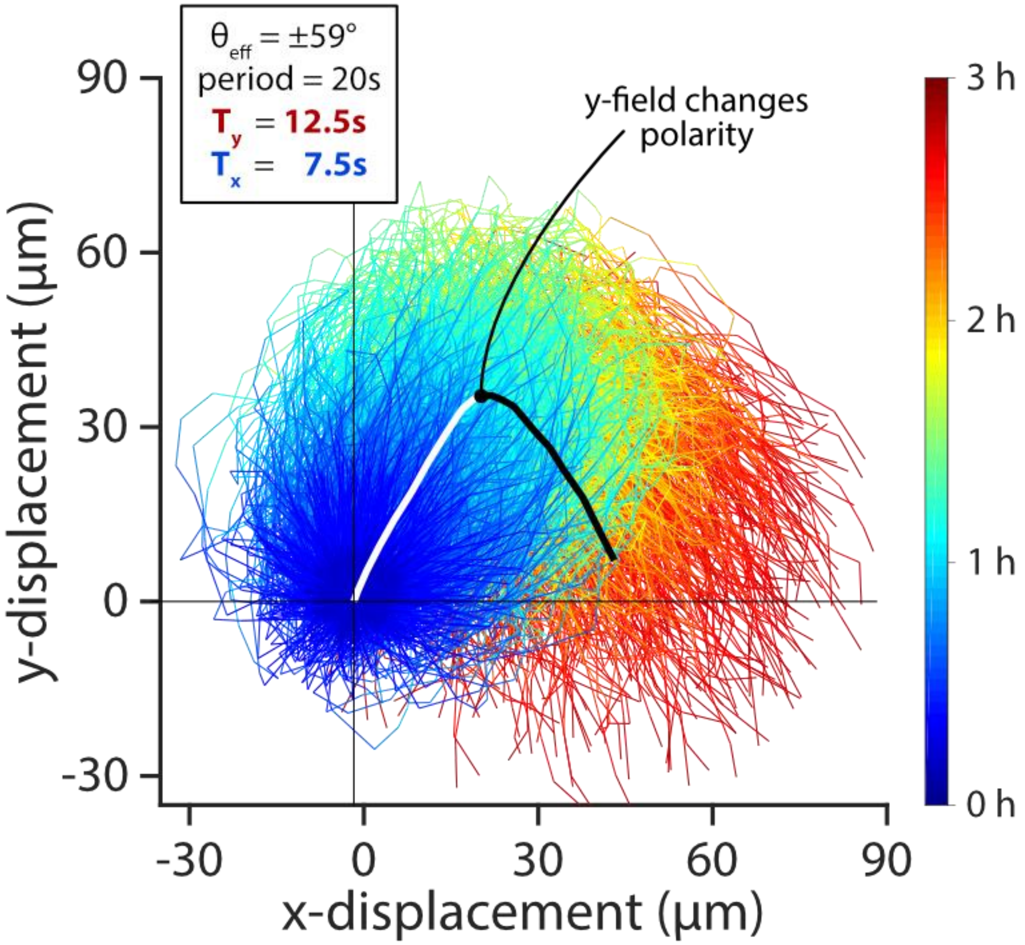


**Supplementary Figure 4:** Trajectories from programmed 60º then -60º diagonals in MDCK monolayer (trajectories measured from n = 2 tissues). Related to Fig. 4. Mean trajectory of individual cell trajectories (white/black). Individual trajectories were taken from 2×2 mm central segments of 5×5 mm tissues during induced migration. For visual clarity, tracks within one standard deviation of the mean cell trajectory are selected and plotted as time-coded multicolor lines (n = 1337 trajectories). See also Supplementary Video 6.

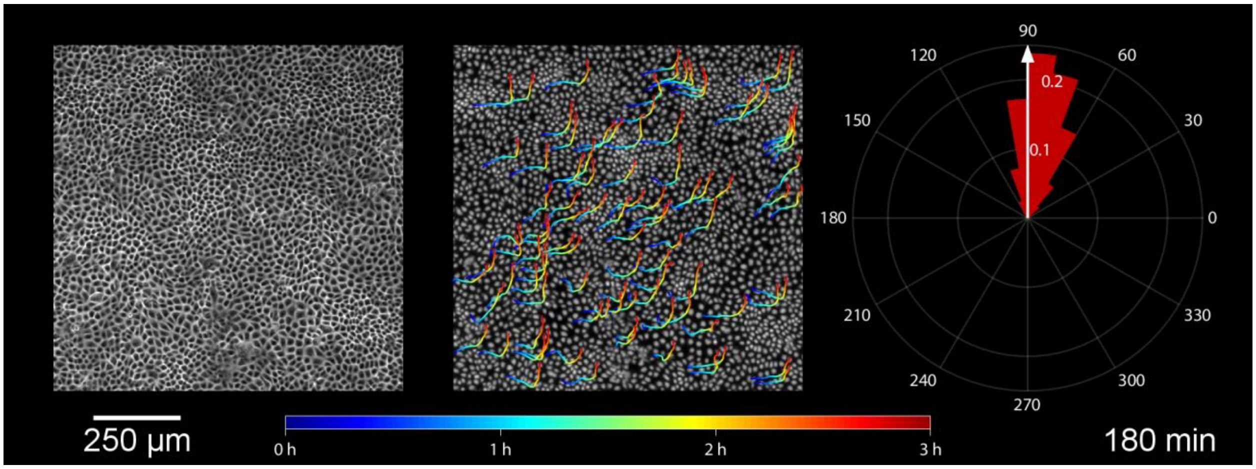


**Supplementary Video 1:** Time-lapse movie of programmed 90º turn in MDCK monolayer. Field was directed for 1.5 h 'right,' then 1.5 h 'up.' Left panel shows phase-contrast microscopy of 1 x 1 mm section in the center of the monolayer. Center panel shows selected trajectories overlaid onto nuclear data labeled by a machine-learning classifier. Right panel shows the field direction (white arrow) overlaid onto a polar histogram of cell displacement angle at each time point. See also Fig. 3.

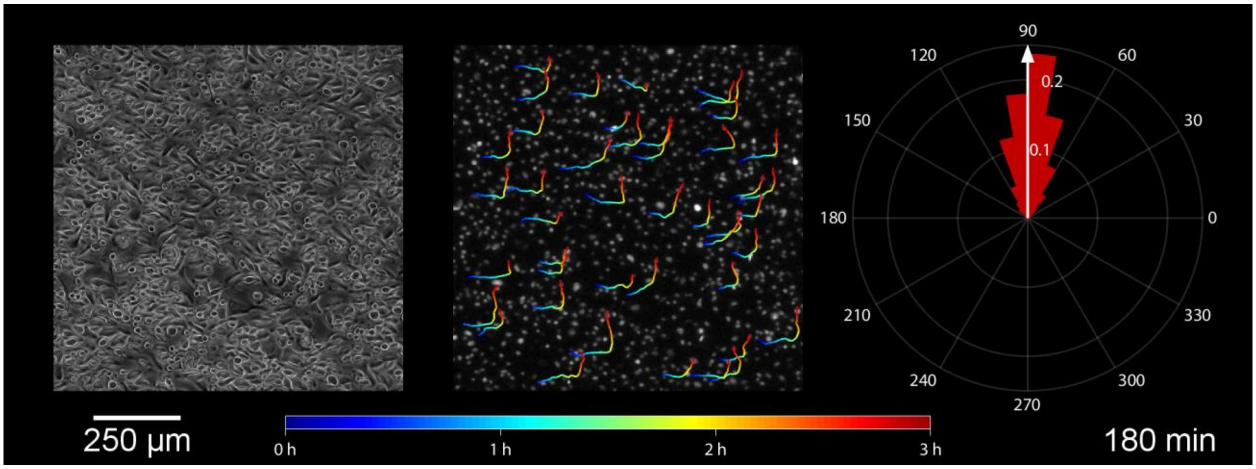


**Supplementary Video 2:** Time-lapse movie of programmed 90º turn in keratinocyte monolayer. Field was directed for 1.5 h 'right,' then 1.5 h 'up.' Left panel shows phase-contrast microscopy of 1 x 1 mm section in the center of the monolayer. Center panel shows selected trajectories overlaid onto nuclear data collected by Hoechst 33342 labeling and fluorescence microscopy. Right panel shows the field direction (white arrow) overlaid onto a polar histogram of cell displacement angle at each time point. See also Fig. 3.

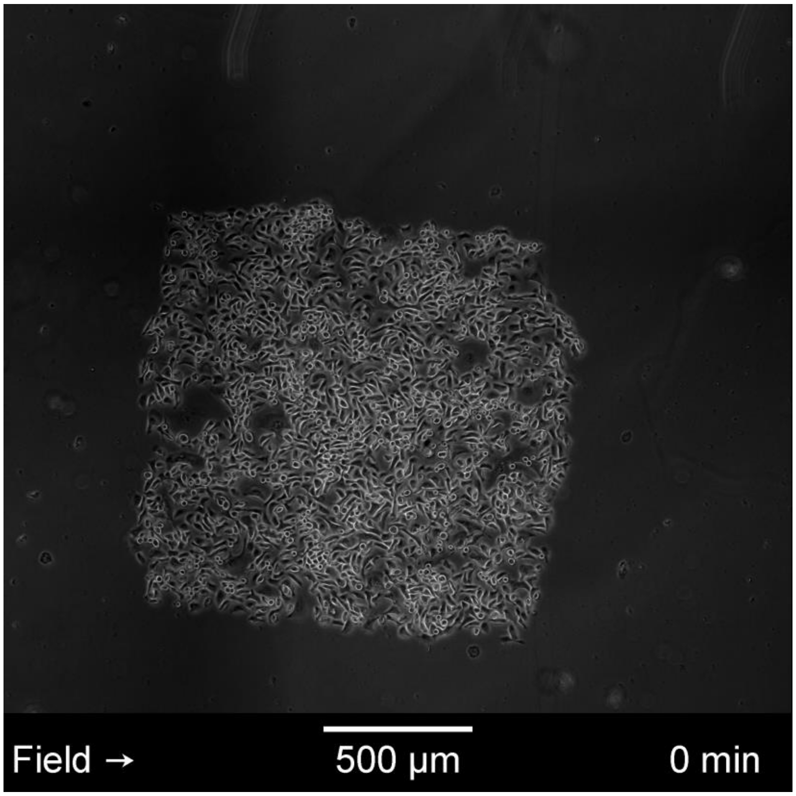


**Supplementary Video 3:** Time-lapse movie of extended 90º turn in keratinocyte monolayer, showing whole monolayer translation. Field was directed for 4 h ‘right,’ then 4 h ‘up.’ See also Fig. 3.

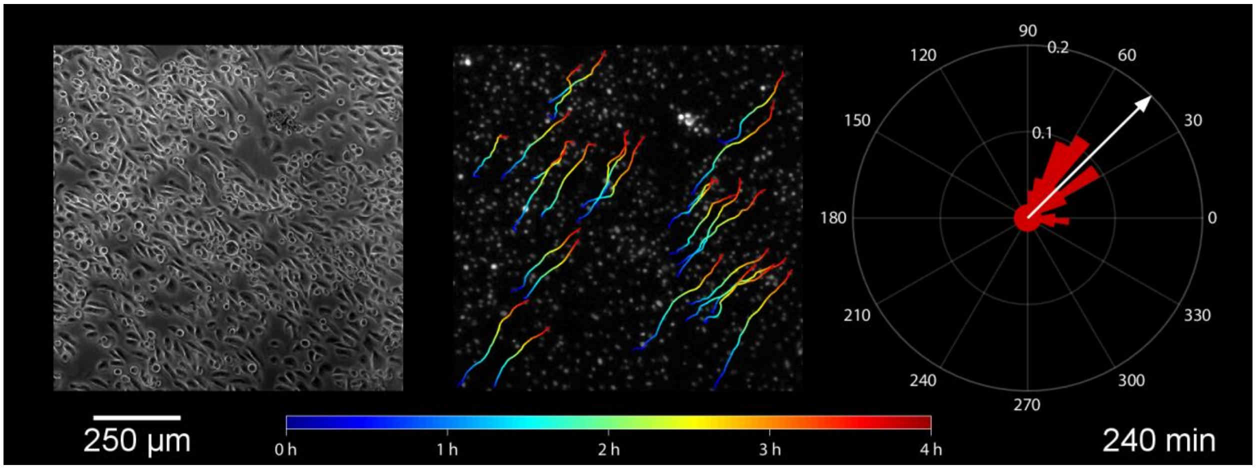


**Supplementary Video 4:** Time-lapse movie of programmed 45º diagonal in keratinocyte monolayer. Field was rapidly alternated between axes for effective stimulation angle of $\theta_{eff}$ = 45º for 4 h. Left panel shows phase-contrast microscopy of 1 x 1 mm section in the center of the monolayer. Center panel shows selected trajectories overlaid onto nuclear data collected by Hoechst 33342 labeling and fluorescence microscopy. Right panel shows the field direction (white arrow) overlaid onto a polar histogram of cell displacement angle at each time point. See also Fig. 4.

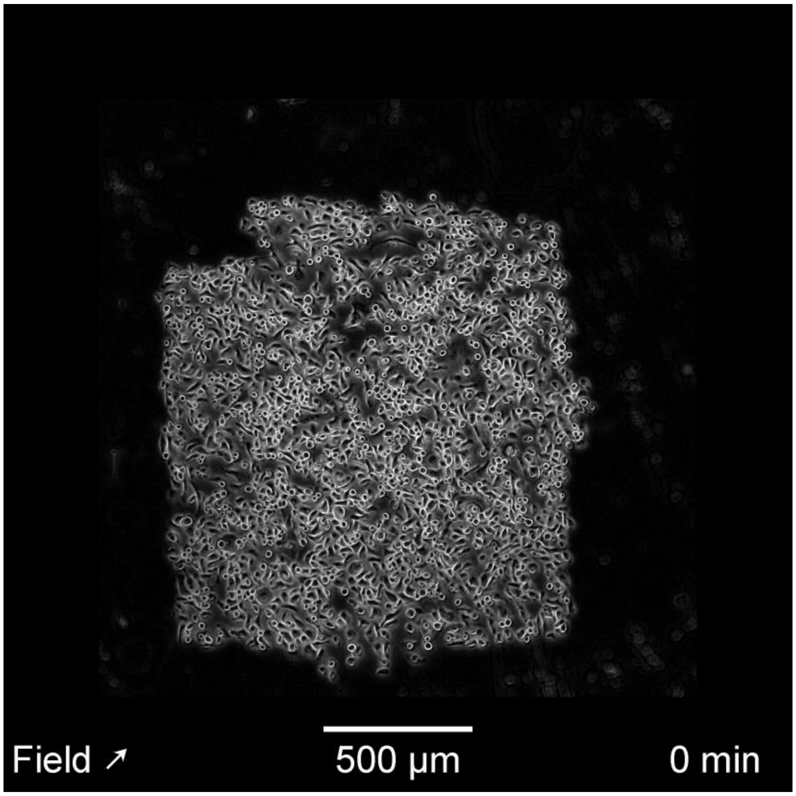


**Supplementary Video 5:** Time-lapse movie of whole tissue translation at 45º angle in keratinocyte monolayer. Field was rapidly alternated between axes for effective stimulation angle of $\theta_{eff}$ = 45º for 4 h. See also Fig. 4.

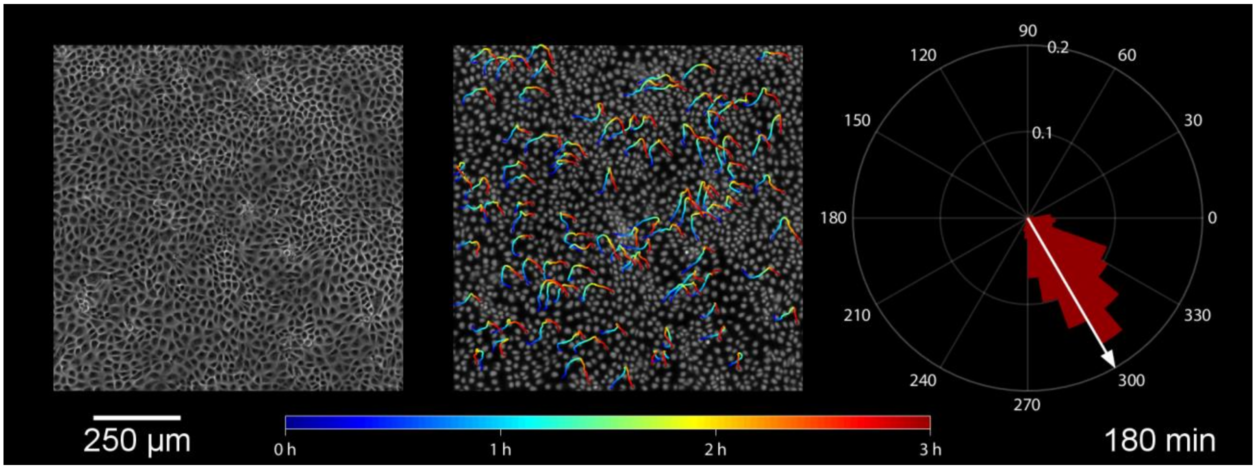


**Supplementary Video 6:** Time-lapse movie of programmed 60º, then -60º diagonals in MDCK monolayer. Field was rapidly alternated between axes for effective stimulation angle of $\theta_{eff}$ = 60º for 1.5 h, then $\theta_{eff}$ = -60º for 1.5 h. Left panel shows phase-contrast microscopy of 1 x 1 mm section in the center of the monolayer. Center panel shows selected trajectories overlaid onto nuclear data labeled by a machine-learning classifier. Right panel shows the field direction (white arrow) overlaid onto a polar histogram of cell displacement angle at each time point. See also Supplementary Fig. 5.

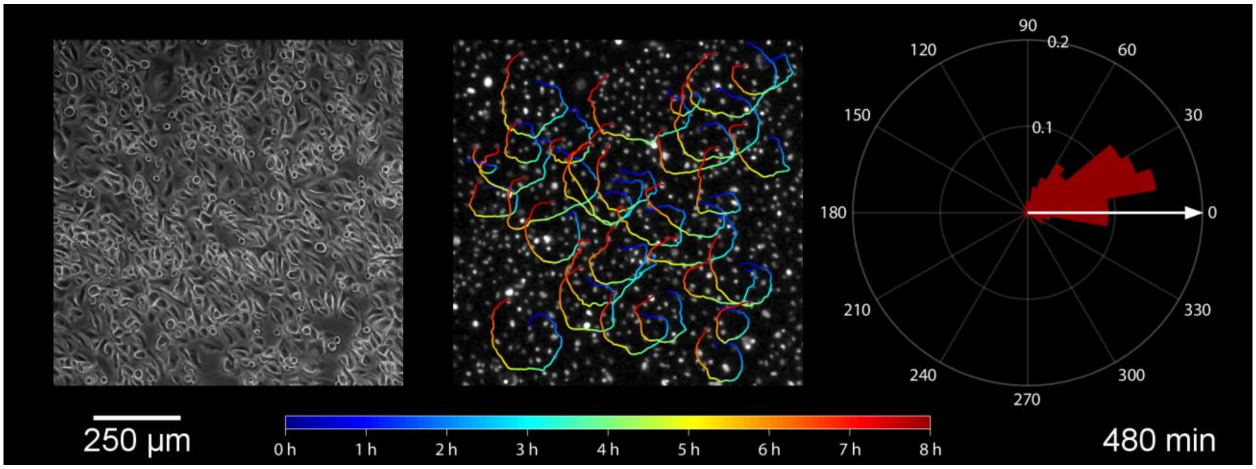


**Supplementary Video 7:** Time-lapse movie of programmed circle maneuver in keratinocyte monolayer. Field was rapidly alternated between axes to rotate the field clockwise at a rate of 45º $h^{-1}$ over 8 h. Left panel shows phase-contrast microscopy of 1 x 1 mm section in the center of the monolayer. Center panel shows selected trajectories overlaid onto nuclear data collected by Hoechst 33342 labeling and fluorescence microscopy. Right panel shows the field direction (white arrow) overlaid onto a polar histogram of cell displacement angle at each time point. See also Fig. 5.
